# Far-Ultraviolet Light Causes Direct DNA Damage in Human Lung Cells and Tissues

**DOI:** 10.1101/2025.03.20.644314

**Authors:** Asta Valanciute, Calum A. Ross, Alex Burden, Syam Mohan P. C. Mohanan, Mohammed Sabbah, Jessica M. W. Ghobrial, Stuart Dickson, Hui Xin Loh, Elvira Williams, Aysha Ali, Christian Brahms, Ahsan R. Akram, Lynn Paterson, Natalie K. Jones, Bethany Mills, John C. Travers, Robert R. Thomson, Kevin Dhaliwal

## Abstract

Far-ultraviolet C (Far-UVC) radiation, with a wavelength range from 200 to 235 nm, is germicidal and holds potential for clinical applications. However, its use against deep-seated and internal infections, such as those affecting the lungs, remains less well established. The safety profile of Far-UVC irradiation requires further investigation across different human tissues.

In this study, we utilised a krypton-chloride (KrCl) excimer lamp and a pulsed laser system to examine the effects of Far-UVC irradiation on human lung cells *in vitro* and primary human tracheal tissue. Primary human tracheal tissue and cells exposed to continuous wave (222 nm) and pulsed 206 nm and 222 nm light at doses of 5, 25, and 50 mJ/cm^2^ exhibited Deoxyribonucleic acid (DNA) damage, including phosphorylation of γH2AX (Ser139). The continuous wave and pulsed 222 nm irradiation caused the formation of pyrimidine-pyrimidone (6-4) photoproducts. Irradiated human lung cells demonstrated reduced viability *in vitro*, and increased lactate dehydrogenase release into the culture medium 48 hours post-irradiation.

Our findings reveal that even low doses of Far-UVC (206 nm, 222 nm) light can penetrate monolayers of human lung epithelial cells, causing direct DNA damage in the form of (6-4) photoproducts and DNA double-strand breaks, ultimately leading to cell death.

## INTRODUCTION

Ultraviolet-C (UVC) light, with wavelengths ranging from approximately 200 nm to 280 nm, is widely recognised for its potent germicidal properties. Recently, the potential for Far-UVC light (200 - 235 nm) to reduce the transmission of airborne pathogens such as SARS-CoV-2 has been extensively studied.^1–4^ At lower doses (6, 17 and 24 mJ/cm^2^), 222 nm light has been shown to inactivate various bacteria, including *Staphylococcus aureus*, *Pseudomonas aeruginosa* and *Escherichia coli*.^5–7^ The germicidal mechanism of 222 nm light is attributed to its ability to degrade aromatic amino acids, oligopeptides, proteases, and proteins, and to damage DNA.^8^

UVC radiation induces direct DNA damage through the formation of a variety of premutagenic and cytotoxic DNA lesions, including pyrimidine dimers. These pyrimidine dimers can be further categorized into cyclobutene pyrimidine dimers (CPDs) and pyrimidine-pyrimidone (6-4) photoproducts ((6-4) PPs). Additionally, UVC radiation absorbed by nucleic acids and proteins can generate oxidative DNA damage, referred to as indirect DNA damage.^9,10^

Despite its highly mutagenic and efficient germicidal activity against viruses and bacteria, 222 nm light has been reported to be non-harmful to mammalian skin and eye cells under certain conditions.^11–16^ However, recent studies have revealed that exposure to 222 nm irradiation can result in significant DNA double-strand breaks (DSBs) in human retinal cell lines (ARPE-19) and HEK-A keratinocytes *in vitro*.^17^ Furthermore, irradiation of monolayered colon epithelial cells with 30 mJ/cm^2^ of 222 nm wavelength was found to compromise cell membrane integrity and reduce cell viability.^18^

The efficacy and safety of Far-UVC light in addressing pulmonary infections, such as ventilator-associated pneumonia caused by multidrug-resistant bacteria or early-stage chronic lung conditions, remain less well-studied. We hypothesised that catheter-delivered Far-UVC irradiation could serve as a potential alternative for treating antibiotic-resistant bacterial infections in the airways. However, no comprehensive studies have been conducted to evaluate the molecular alterations induced by Far-UVC irradiation in human lung epithelial cells or lung tissue. Furthermore, the biological effects of exposure to low doses of continuous wave (CW) or pulsed Far-UVC irradiation on human lung epithelial cells remain unexplored.

The primary objective of this study was to investigate the biological effects of 206 nm and 222 nm irradiation on DNA damage in human lung cells, with the aim of assessing its safety and potential as a tool for a wide range of clinical applications.

## METHOD DETAILS

All chemicals, cell lines, media, antibodies, fluorescence probes, commercial assays and tools are included in Key resources table, Supplementary Material.

### Experimental sources of Far-UVC irradiation

In this study, we used two Far-UVC light sources to assess the effects of irradiation on human cells *in vitro* and tracheal tissue. The first source was a krypton-chloride (KrCl) excimer lamp (Ushio Care222 module; Key resources table, Supplementary Material), a CW system equipped with filters to maintain a constant 222 nm output. This lamp was positioned 20 cm from the sample to ensure uniform UV intensity across an entire six-well plate, a setup used consistently in all experiments.

The second light source has been described by Brahms et al.^19^ It is an ultrafast pulsed laser system based on frequency conversion in gas-filled hollow capillary fibres, capable of tuning from 206 nm to 254 nm wavelengths. In this study, pulsed 254 nm irradiation served as a positive control for DNA damage induction in mammalian cells.^20^ The pulsed source operated at a 10 kHz repetition rate, with its output beam expanded and collimated up to a diameter of ~20 mm before being directed onto individual samples in well plates.

To compare the impact of CW and pulsed light, the intensity of both sources at the sample was measured and adjusted to be approximately 0.28 mW/cm^2^. Total irradiation doses of 5, 25, and 50 mJ/cm^2^ were achieved by varying exposure times. Ozone generation from both CW and pulsed Far-UVC sources was considered minimal and insignificant due to the short exposure duration.

### Cell culture

Human lung cell lines, including Beas-2B (immortalised human bronchial epithelial cells), H1299 (human non-small cell lung carcinoma cells), and HaCaT (human keratinocytes), were obtained from ATCC (Manassas, VA, USA). Cells were cultured in complete medium consisting of DMEM supplemented with 10% fetal bovine serum (FBS), 100 IU/mL penicillin G sodium salt, 100 µg/mL streptomycin sulfate, and 2 mM L-glutamine. They were routinely passaged in T25 cell culture flasks upon reaching 80– 90% confluence. Mycoplasma contamination was regularly tested, and all assays were conducted using mycoplasma-free cell cultures.

### Human trachea

Human lungs that were unsuitable for transplantation were obtained from the National Health Service Blood and Transplant (NHSBT). The studies involving this human tissue were approved by the London – Central Research Ethics Committee (REC No: 16/LO/1883) and supported by NHS Lothian SAHSC Bioresource (REC No: 20/ES/0061).

Trachea was excised from the donated lungs and washed with 50 mL of PBS solution and then cut into 1 cm^2^ pieces. Five hundred microliters of DMEM complete medium (without phenol red) were applied to the tissue. The trachea, with its airway ciliated cells exposed, was then directly irradiated with Far-UVC light.

### Immunofluorescence for human cells

Cells were seeded on coverslips and cultured for 24 hours before exposure to 222 nm light, followed by an additional 2-hour incubation. Cells were then fixed in 4% paraformaldehyde for 15 minutes and incubated in a permeabilization/blocking buffer (1% BSA, 10% FBS, 0.1% Triton X-100, 0.1% Saponin) for 1 hour at room temperature (RT).

Primary antibody for γH2AX(Ser139) was diluted (1:200) and incubated overnight at 4°C, followed by a 1-hour incubation with secondary antibody (Alexa Fluor^TM^488 anti-Rabbit IgG, 1:1000) and Hoechst 33342 (1:1000) at RT in the dark. Samples were then mounted and imaged using a Leica SP8 confocal microscope with a 40x objective.

Fluorescence intensity was quantified using ImageJ software, with corrected total cell fluorescence (CTCF) calculated as: Integrated Density / (Cell Area × Mean Background Fluorescence).

Etoposide, a positive control (a final concentration of 50 µM, Etop), was used to induce DNA DSBs.

### Immunofluorescence of formalin-fixed paraffin-embedded human trachea

Irradiated-human trachea samples and controls were fixed in 4% neutral buffered formalin overnight at RT, processed using a standard automated tissue processor, and embedded in paraffin wax. Sections (5–8 µm) were cut for labelling experiments.

Slides were dewaxed sequentially in xylene, ethanol gradients (100%, 90%, 70%, 40%), and dH₂O, then transferred to citrate buffer (pH 6.0) for antigen retrieval via microwave heating (20 minutes). After cooling (30 minutes) and rinsing in running water (15 minutes), slides were placed in PBS to prevent drying. Tissue sections were framed with a PAP pen and blocked overnight at 4°C in PBS containing 1% BSA, 10% FBS, 0.1% Triton X-100, and 0.3% Saponin.

After washing with PBS, slides were incubated with primary antibody: anti-γH2AX(Ser139) or anti-(6-4) DNA photoproducts in immunofluorescence buffer (PBS with 3% BSA, 1% goat serum, 0.1% Triton X-100, 0.1% Saponin) overnight at 4°C. Each primary antibody was incubated with a different tissue cut. A small parafilm cover was used to prevent drying. After washing with 0.5% Tween 20 in PBS, slides were incubated with secondary antibody (Alexa Fluor^TM^488 anti-Rabbit IgG or Alexa Fluor^TM^594 anti-Rabbit IgG, 1:1000, respectively) and Hoechst 33342 (1:1000) for 1 hour at RT in the dark.

Following washes with 0.5% Tween 20 in PBS, Vector TrueVIEW reagent was applied (5 min) to reduce autofluorescence. Slides were mounted with Vectashield Vibrance Antifade Mounting Medium and imaged using a Leica SP8 confocal microscope with a 40x objective. Image reconstruction was performed with ImageJ software.

### Annexin V apoptosis detection

Irradiated cells and controls were incubated in a humidified atmosphere at 37°C with 5% CO_2_ for 12 hours. The medium was aspirated and preserved for the identification of apoptotic cells. The adherent cells were washed with 2 mL of PBS. After washing, cells were incubated with 0.5 mL of TrypLE Express/without Phenol Red for one minute at RT. After adding 2 mL of PBS, a cell suspension, including non-adherent and adherent cells, was centrifugated at 400 g for 5 minutes. For each treatment 100 µL of cell suspension (1 x 10^6^ cells) was mixed with FITC Annexin V (2.5 µL) and Propidium Iodide Solution (1 µL, PI), a viability dye, and incubated for 15 minutes at RT protected from light. The cells were washed with 400 µL of Annexin V Binding Buffer and centrifugated at 400 g for 5 minutes. The cells were resuspended in 250 µL of Annexin V Binding Buffer and analyzed by Cytek Aurora (Cytek Biosciences). The flow cytometry data was analyzed by FlowJo software. Staurosporine, a positive control (a final concentration of 1.25 µM, ST), was used to induce a non-lytic apoptosis.

### Crystal violet assay

To assess the effect of Far-UVC exposure on cell viability, cells were seeded into six-well plates at 0.35 × 10⁶ cells per well the day before irradiation, reaching 70–80% confluence at the time of irradiation. Cells were incubated in 500 µL of phenol red-free DMEM before being exposed to a specific Far-UVC dose and wavelength. After irradiation, the medium was replaced with fresh DMEM (2 mL per well), and cells were incubated at 37°C with 5% CO₂ for the required duration.

Cells were then washed twice with PBS, fixed in 4% paraformaldehyde for 15 minutes at RT, and stained with 1 mL of 0.5% crystal violet (CV) in 20% methanol for 20 minutes at RT. After staining, cells were washed four times with dH₂O, air-dried, and solubilized in 1 mL of 30% acetic acid solution. The plate was incubated for 20 minutes with occasional shaking. Optical density (OD) was measured at 570 nm using a Synergy H1 Hybrid Reader (BioTek Instruments, Winooski, VT, USA). Etoposide (a final concentration of 50 µM, Etop) was used as a positive control.

### Lactate dehydrogenase (LDH) cytotoxicity assay

Cellular LDH released into the cellular medium upon plasma membrane damage by 222 nm light was assessed with CyQuant LDH Cytotoxicity assay kit according to the manufacturer’s instructions. The absorbance of LDH (OD) was measured at 490 nm using a Synergy H1 Hybrid Reader (BioTek Instruments, Winooski, VT, USA). Etoposide (a final concentration of 50 µM, Etop) was used as a positive control.

### Statistical analysis

Statistical analysis was carried out using GraphPad Prism10. Statistical differences were determined via one-way ANOVA with comparison based on a control column. Independent experiments were performed at least three times or otherwise as stated. Results are expressed as the mean ± SD. p-value < 0.05 was considered significant. *p < 0.05, **p <0.01, ***p <0.001, ****p < 0.0001, respectively.

## RESULTS

### 222 nm-induced DNA DSBs in monolayered human cells

Previous studies have shown that UV-B (280–315 nm)^10,20^ and Far-UVC wavelengths: 206 nm^21^ and 222 nm^17^ can penetrate cells, traverse the cytoplasm, reach the nucleus, and induce DNA damage, leading to an excess of DNA double-strand breaks.

We investigated DNA DSBs damage in human cells, irradiated with CW 222 nm light at doses of 5 and 25 mJ/cm^2^, using immunofluorescence staining for γH2AX(Ser139) 2 hours after irradiation. Irradiated Beas-2B cells exhibited a dose-dependent increase in DNA DSBs, as evidenced by γH2AX(Ser139) staining. Hoechst staining revealed no nuclear aggregation or fragmentation in the irradiated Beas-2B cells (Figure 1A). Furthermore, CW 222 nm exposure resulted in significantly higher γH2AX(Ser139) fluorescent intensity compared to non-irradiated controls (Figure 1B).

**Figure 1.**
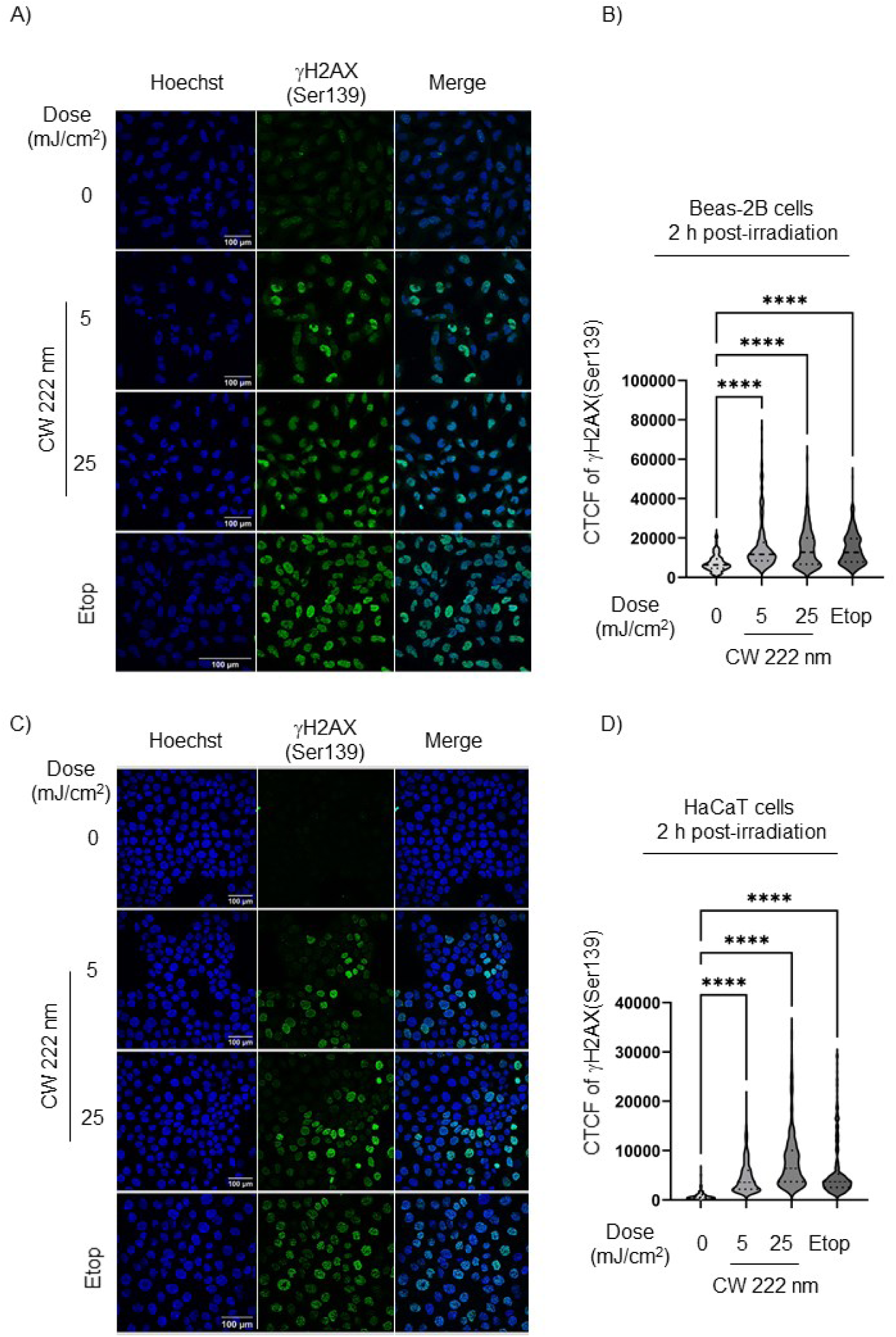
CW 222 nm-induced DNA DSBs damage in monolayered Beas-2B and HaCaT cells *in vitro*. (A) Representative immunofluorescence images of γH2AX(Ser139) staining (green) in Beas-2B cells 2 hours post-irradiation with CW 222 nm. Hoechst nuclei (blue). An untreated sample (0 mJ/cm^2^) was used as an internal control. Etop was used as a positive control of γH2AX(Ser139) staining. Scale bar, 100 μm. (B) Quantification of the γH2AX(Ser139) staining as the corrected total cell fluorescence (CTCF) was performed on Beas-2B cells 2 hours post-irradiation. Data presented as means +/−SD from n = 3 experiments. In total 306 cells were analyzed per condition. One-way ANOVA with repeated measures (****p < 0.0001). (C) Representative immunofluorescence images of γH2AX(Ser139) staining (green) in HaCaT cells at 2 hours following CW 222 nm irradiation. Hoechst nuclei (blue). Scale bar, 100 μm. (D) Quantification of the γH2AX(Ser139) activation was performed on HaCaT cells 2 hours post-irradiation with CW 222 nm. Data presented as means +/−SD from n = 3 experiments. In total 361 cells were analyzed per condition. One-way ANOVA with repeated measures (****p < 0.0001).

Similarly, CW 222 nm irradiation increased γH2AX(Ser139) phosphorylation in HaCaT keratinocytes (Figure 1C, D), a skin cell model known for its resistance to Far-UVC light exposure.

We further evaluated the DNA damage response in pulsed 222 nm- and 206 nm-irradiated Beas-2B cells *in vitro*. Following 5 and 25 mJ/cm^2^ of pulsed 222 nm and 206 nm irradiation, monolayer-cultured Beas-2B cells showed a dose-dependent increase in γH2AX(Ser139) phosphorylation compared to non-irradiated cells (Figure 2A, B, C, D, respectively).

**Figure 2.**
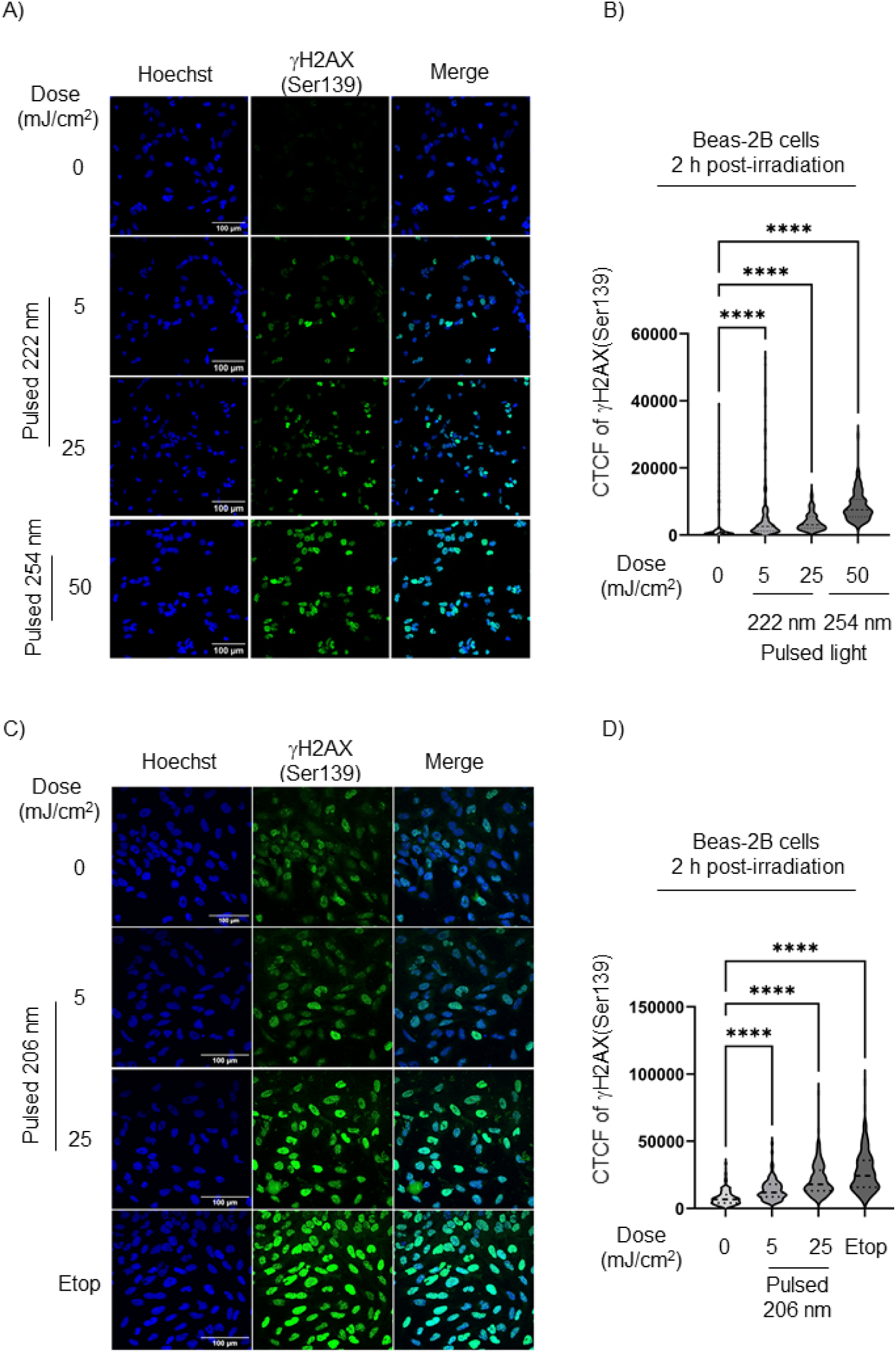
Pulsed 222 nm and 206 nm irradiation induced DNA DSBs damage in monolayered Beas-2B *in vitro*. (A) Representative immunofluorescence images of γH2AX(Ser139) staining (green) in Beas-2B cells 2 hours post-irradiation with pulsed 222 nm. Hoechst nuclei (blue). Pulsed 254 nm light (50 mJ/cm^2^) was used as a positive control of γH2AX(Ser139) staining. Scale bar, 100 μm. (B) Quantification of the γH2AX(Ser139) staining as the corrected total cell fluorescence (CTCF) was performed on Beas-2B cells 2 hours post-irradiation. Data presented as means +/−SD from n = 3 experiments. In total 237 cells were analyzed per condition. One-way ANOVA with repeated measures (****p < 0.0001). (C) Representative immunofluorescence images of γH2AX(Ser139) staining (green) in Beas-2B cells at 2 hours following pulsed 206 nm irradiation. Hoechst nuclei (blue). Scale bar, 100 μm. (D) Quantification of the γH2AX(Ser139) activation was performed on Beas-2B cells 2 hours post-irradiation with pulsed 206 nm. Data presented as means +/−SD from n = 3 experiments. In total 232 cells were analyzed per condition. One-way ANOVA with repeated measures (****p < 0.0001).

### 222 nm-induced DNA double-strand breaks and (6-4) PPs in human trachea

Building on our findings of 222 nm-induced DNA DSBs in monolayer-cultured cells 2 hours after irradiation, we hypothesised that DNA in the ciliated pseudostratified columnar epithelium of the human trachea could also be damaged by 222 nm irradiation.

Using fluorescent confocal microscopy and γH2AX(Ser139) staining, we observed that CW 222 nm, pulsed 222 nm and pulsed 206 nm irradiation, at doses of 5, 25, and 50 mJ/cm^2^, resulted in a significantly higher number of γH2AX(Ser139)-positive cells compared to non-irradiated tissue 10 minutes after the initial irradiation (Figure 3A, B, Figure S1A, B, respectively).

**Figure 3.**
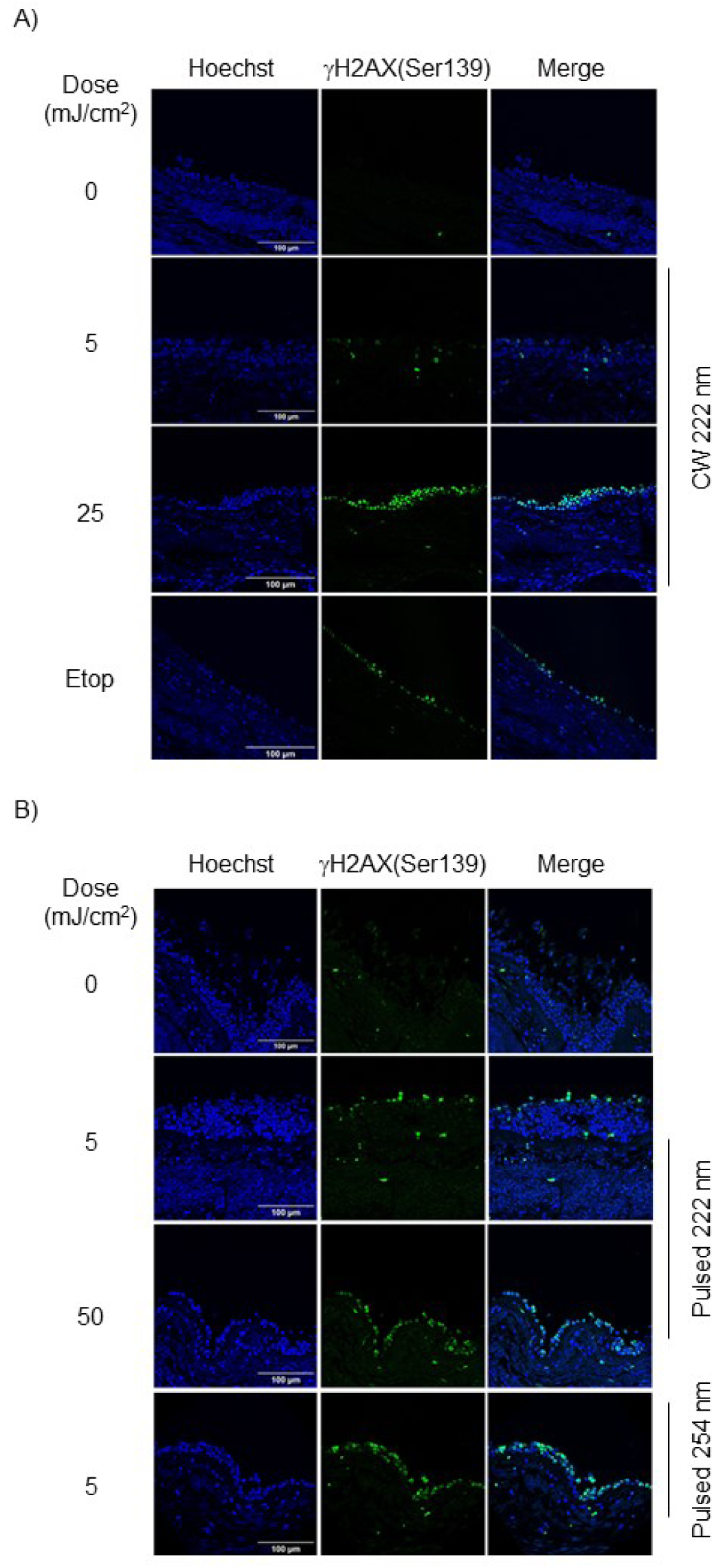
CW and pulsed 222 nm irradiation induced DNA DSBs in human trachea. (A) Representative immunofluorescence images of γH2AX(Ser139) staining (green) in human trachea after CW 222 nm irradiation. Hoechst nuclei (blue). Scale bar, 100 μm. Representative immunofluorescence images of γH2AX(Ser139) staining (green) after pulsed 222 nm irradiation in human trachea. Hoechst nuclei (blue). Pulsed 254 nm (5 mJ/cm^2^) irradiation was used as a positive control. Scale bar, 100 μm.

In addition to causing DNA DSBs, UVC radiation also generates thousands of DNA lesions in living cells,^22,23^ for example, (6-4) PPs. We investigated the formation of (6-4) PPs in human tracheal tissue following CW and pulsed 222 nm irradiation at energy doses of 5, 25, and 50 mJ/cm^2^. Both CW and pulsed 222 nm irradiation induced (6-4) PPs formation in the upper epithelial cells of the trachea in a dose-dependent manner, observed 10 minutes after irradiation (Figure 4A, B, and Figure S2).

**Figure 4.**
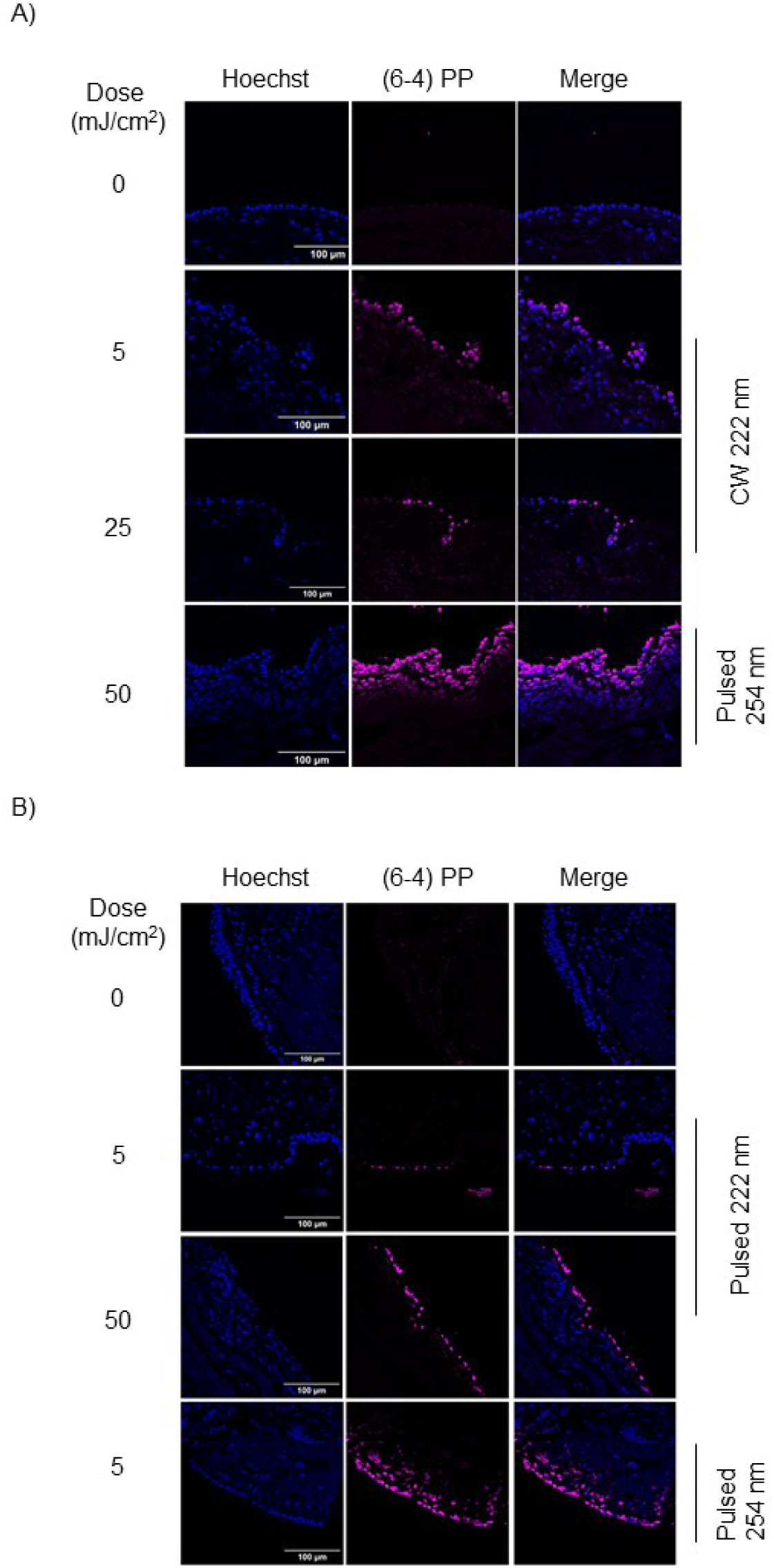
(6-4) PP-expressing cells were detected in human trachea after 222 nm irradiation. (A) Representative immunofluorescence images of (6-4) PP staining (magenta) in human trachea 10 minutes after CW 222 nm irradiation. Hoechst nuclei (blue). Pulsed 254 nm-irradiated samples were used as a positive control. Scale bar, 100 μm. (B) Representative immunofluorescence images of (6-4) PP staining (magenta) in human trachea 10 minutes after exposure to pulsed 222 nm light. Hoechst nuclei (blue). Scale bar, 100 μm.

### Impact of 222 nm light on cell viability and apoptosis in human cells *in vitro*

After identifying significant DNA damage in human lung cells exposed to 222 nm light, we explored cell survival 48 hours post-irradiation. After CV staining, we localized the irradiation spot in the human adherent cells *in vitro*, also we observed that two human lung cell lines (Beas-2B and H1299) and HaCaT keratinocytes subjected to CW 222 nm irradiation failed to proliferate within 48 hours of exposure. At this time point, a distinct irradiated area with reduced CV staining intensity (highlighted by a yellow square bracket) was evident in all three CW 222 nm-irradiated cell lines (Figure 5A). Optical density measurements of CV at 570 nm confirmed that CW 222 nm irradiation induced cytotoxicity in a dose-dependent manner across all three cell lines. (Figures 5B, C, and D, respectively).

**Figure 5.**
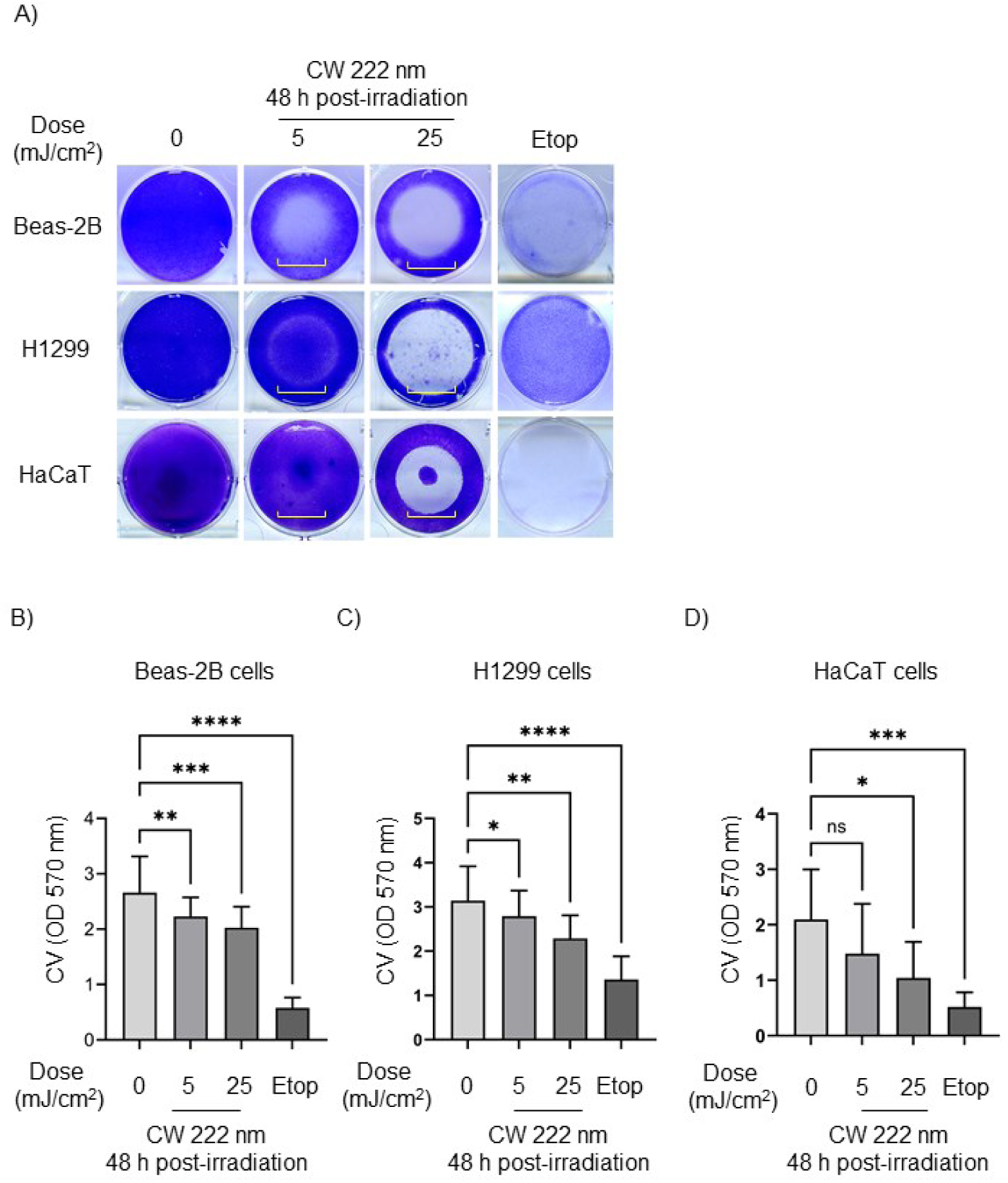
CW 222 nm irradiation resulted in decreased cell viability in human cells *in vitro* 48 hours post-irradiation. (A) Representative images of CV staining on Beas-2B, H1299 and HaCaT cells 48 hours after CW 222 nm irradiation. A square yellow bracket indicates an irradiated area. (B) The OD of CV-stained Beas-2B cells was measured at 570 nm. Data are shown as mean +/− SD from n = 3 experiments. (C) The OD of CV-stained H1299 was measured at 570 nm. Data are presented as mean +/− SD from n = 3 experiments (D) The OD of CV-stained HaCaT cells was measured at 570 nm Data are presented as mean +/− SD from n = 3 experiments. Where applicable, one-way ANOVA with repeated measures was performed (ns, not significant, *p < 0.05, **p < 0.01, ***p <0.001, ****p < 0.0001.

After identifying a decreased proliferation and survival of CW 222 nm-irradiated cells 48 hours post-irradiation, we next interrogated whether CW 222 nm-irradiated cells show apoptosis at earlier timepoint (12 hours post-irradiation). Using FITC Annexin V and PI staining with flow cytometry, we found that Beas-2B cells irradiated with CW 222 nm light showed reduced viability compared to non-irradiated cells 12 hours post-irradiation (Figure 6A). Specifically, CW 222 nm irradiation at doses of 25 and 50 mJ/cm^2^ resulted in a significantly decrease percentage of live cells (36% and 30%, respectively) compared to non-irradiated cells (82%) (Figure 6A). This reduction correlated with increased late apoptosis, as indicated by positive Annexin V and PI staining (Figure 6B).

**Figure 6.**
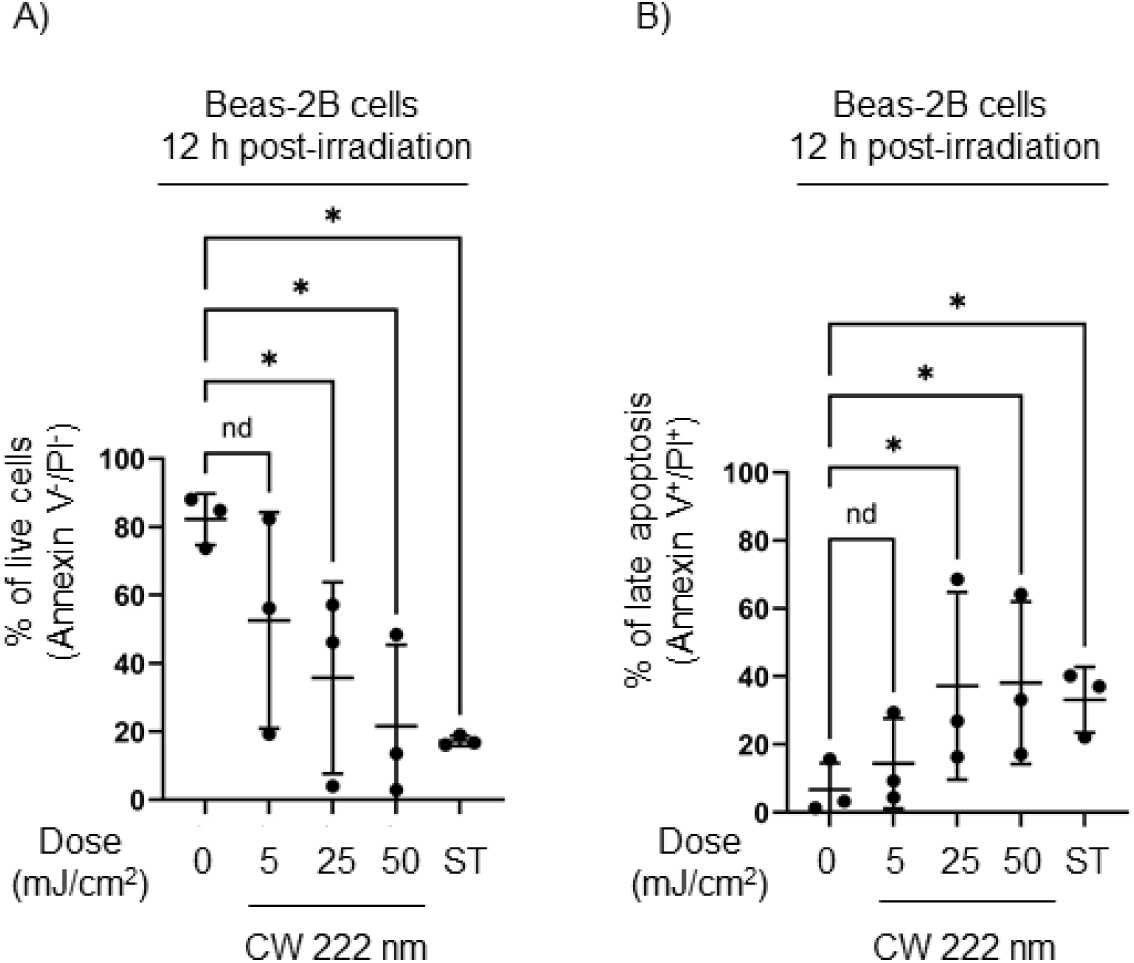
CW 222 nm irradiation reduced cell viability and induced apoptosis 12 hours post-irradiation in Beas-2B cells. (A) Decreased Beas-2B cell viability was demonstrated by FITC Annexin V and PI assay via flow cytometry 12 hours post-irradiation. Data presented as means +/− SEM from n = 3 experiments with representative flow cytometry dot plots per treated group. (B) Late apoptosis demonstrated in Beas-2B cells 12 hours post-irradiation by Annexin V assay. Data presented as means +/− SEM from n = 3 experiments with representative flow cytometry dot plots per treated group. Were applicable, data (A) and (B) was analysed using the Friedman test (nd, not a discovery, *p < 0.05).

After determining the cell-killing effect of low-dose CW 222 nm irradiation *in vitro*, we next evaluated whether both pulsed 222 nm and 206 nm lights could reduce cell viability 48 hours post-irradiation. Pulsed 222 nm irradiation decreased the viability of Beas-2B cells, as indicated by reduced CV staining (highlighted by a yellow square bracket) at 48 hours post-exposure. Beas-2B cells exposed to pulsed 222 nm irradiation (5, 25 mJ/cm^2^) did not survive and detached from the plate, whereas non-irradiated cells remained viable (Figure 7A). Optical density measurements of CV staining confirmed the cytotoxic effects of pulsed 222 nm irradiation in a dose-dependent manner, inhibiting the expansion of irradiated cells 48 hours post-exposure (Figure 7B). In concordance with CW 222 nm and pulsed 222 nm irradiation, pulsed 206 nm irradiation also decreased viability of Beas-2B cells 48 hours post-irradiation (Figure 7C, D).

**Figure 7.**
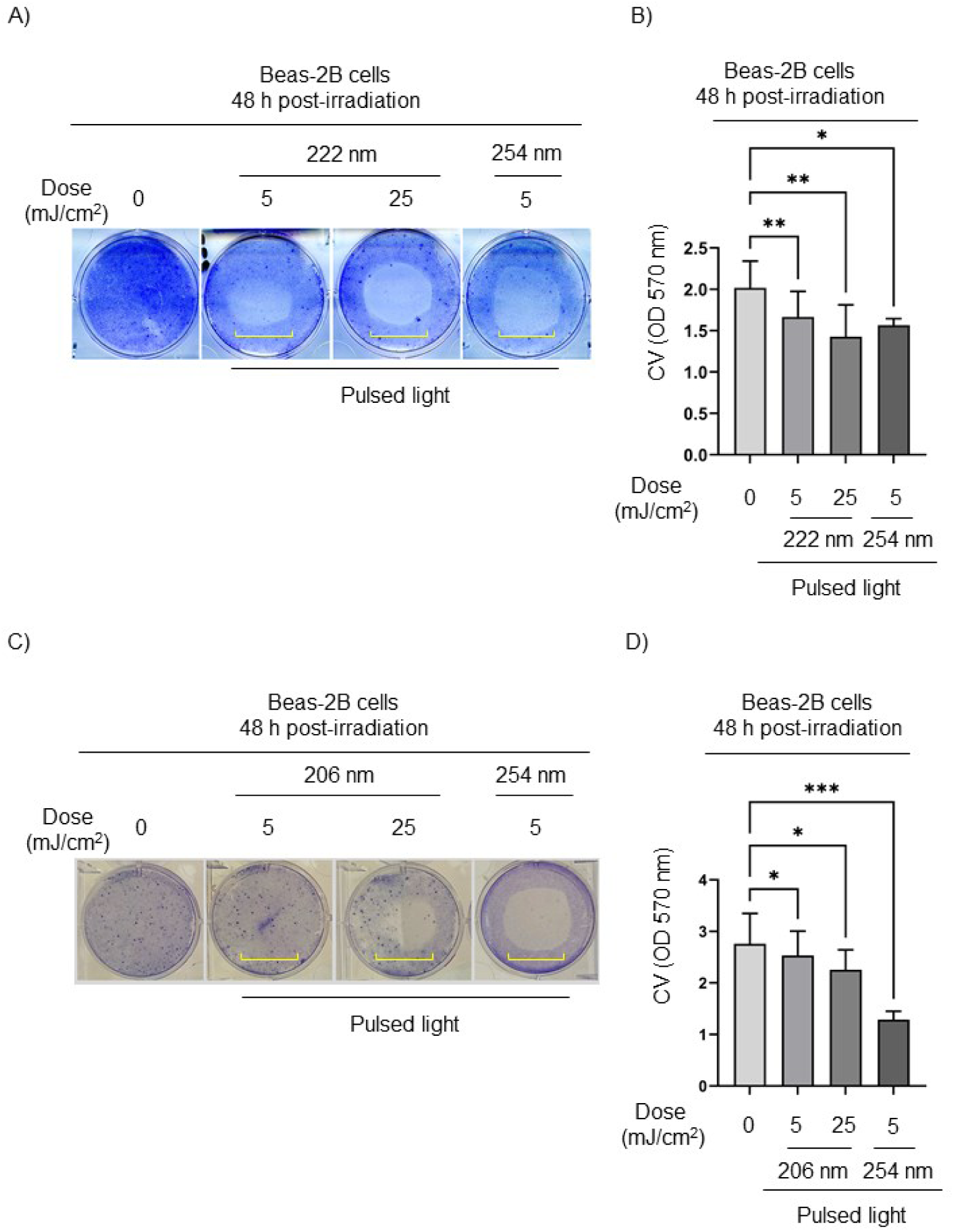
Decrease of CV staining in pulsed 222 nm- and 206 nm-irradiated Beas-2B cells. (A) Representative images of CV staining on Beas-2B cells at 48 hours after pulsed 222 nm irradiation. A square yellow bracket indicates an irradiated area. (B) The OD of CV-stained Beas-2B cells 48 hours post-irradiation with pulsed 222 nm light. Data are shown as mean +/− SD from n = 3 experiments (C) Representative images of CV-stained Beas-2B cells 48 hours after pulsed 206 nm irradiation. A square yellow bracket indicates an irradiated area. (D) The OD of CV-stained-Beas-2B cells 48 hours post-irradiation with pulsed 206 nm light. Data are shown as mean +/− SD from n = 3 experiments. Where applicable, one-way ANOVA with repeated measures was performed (*p < 0.05, **p < 0.01, ***p < 0.001).

It has also been shown that UVC irradiation can damage the cell membrane leading the cell content to leak into the cell medium.^24^ A lactate dehydrogenase cytotoxicity assay was performed to investigate possible cell membrane damage after CW 222 nm irradiation. We found that CW 222 nm irradiation damaged the cell membrane in Beas-2B and H1299 cells 48 hours post-irradiation at a dose of 25 mJ/cm^2^ compared to non-irradiated cells as indicated by an increase in LDH release into the culture medium (Figure 8A, B, respectively).

**Figure 8.**
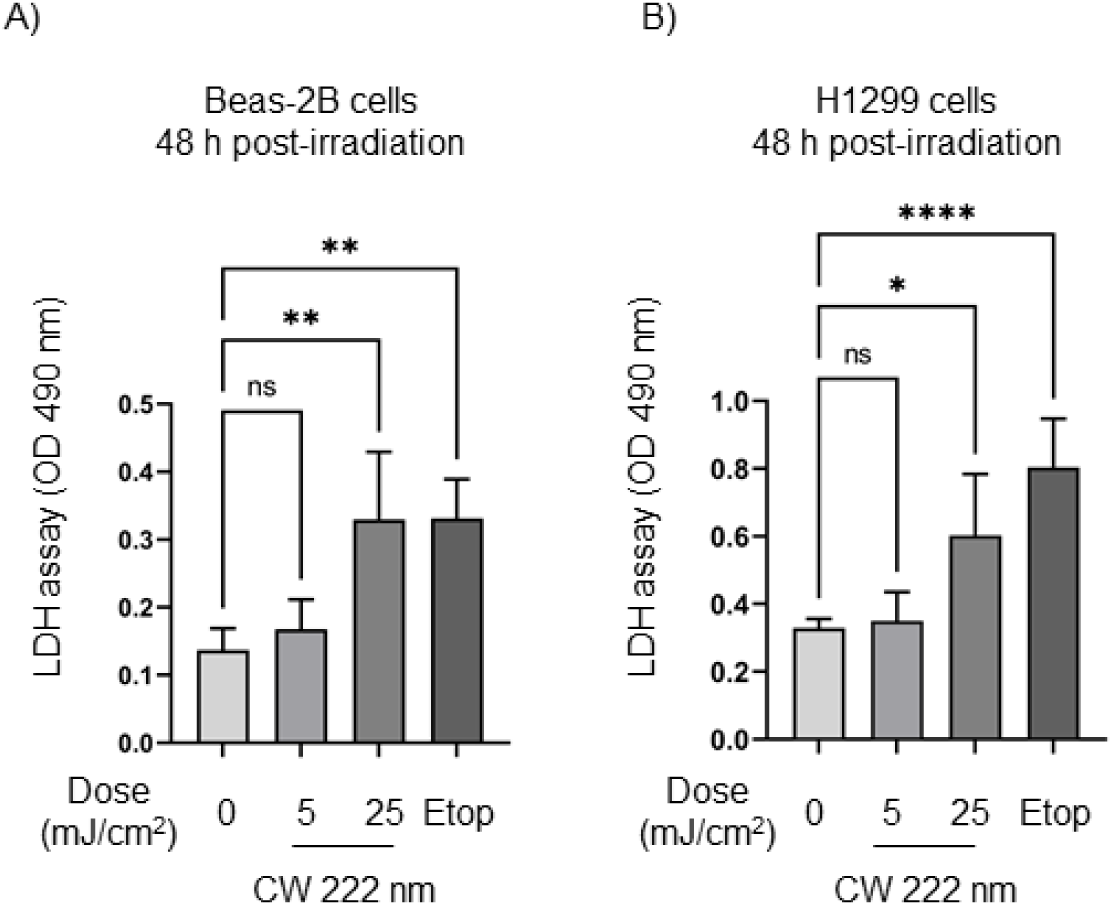
CW 222 nm irradiation induced an increase of lactate dehydrogenase into cell culture media 48 hours post-irradiation. The extracellular LDH levels in the culture media from CW 222 nm-irradiated (A) Beas-2B and (B) H1299 cells measured at 490 nm. Data presented as means +/−SD from n = 3 experiments. One-way ANOVA with repeated measures (ns, not significant, *p < 0.05, **p < 0.01, ****p < 0.0001).

## DISCUSSION

Far-UVC light with a wavelength of 222 nm has been reported as an effective antimicrobial and antiviral agent.^1,2,5^ In recent years, multiple studies have investigated its impact on skin models, concluding that 222 nm irradiation offers high germicidal efficacy with minimal safety concerns compared to longer UVC wavelengths such as 254 nm.^11,25,26^ If 222 nm light can inactivate antibiotic-resistant bacteria while being well-tolerated by human cells without inducing DNA damage or photocarcinogenesis, it could be a promising option for clinical applications to prevent colonisation and treat antibiotic-resistant pathogens in the airways. However, a key issue remains determining the safety of Far-UVC irradiation on human lung cells and tissues.

Recent studies have shown that CW 222 nm irradiation at a dose of 87.6 mJ/cm^2^ increased the percentage of γH2AX-positive cells and reduced proliferation in human retinal pigment epithelial cells compared to non-irradiated controls.^17^ In our study, we found that CW 222 nm, pulsed 222 nm and pulsed 206 nm irradiation (5, 25, 50 mJ/cm^2^) induced γH2AX phosphorylation at Ser139, a marker of DNA DSBs, in a dose-dependent manner in human lung cells and in human tracheal tissue.

Previous studies have demonstrated that 254 nm UV irradiation causes DNA lesions.^12,15,20^ Unlike 254 nm, 222 nm irradiation has been assumed safe for skin and eye models without inducing CPDs or (6-4) PPs.^12,13,27^ However, 311 nm irradiation (30 mJ/cm^2^) has been shown to generate (6-4) PPs in human skin fibroblasts, with (6-4) PPs — rather than CPDs — playing a critical role in impeding DNA replication and leading to cell death.^28^ In our study, we detected a strong upregulation of (6-4) PPs in the human tracheal epithelium after CW and pulsed 222 nm exposure at low doses (5, 25, 50 mJ/cm^2^).

Prolonged, unrepaired DNA lesions and DNA DSBs following UV exposure can impair cell growth and lead to cell death. Both Ong et al.^17^ and Nishikawa et al.^18^ reported that 222 nm-irradiated cells exhibited increased apoptosis and reduced viability. In our study, we observed a significant decrease in cell viability 12- and 48-hours post-irradiation, with no recovery, indicating continuous cell death beyond initial exposure.

We demonstrated both the short (10 minutes or 2 hours) and long-term (12 and 48 hours) effects of Far-UVC irradiation on human lung cells and tracheal tissue. A single low-dose exposure (5, 25, 50 mJ/cm^2^) of CW 222 nm, pulsed 222 nm and pulsed 206 nm irradiation caused direct DNA damage in the nucleus, leading to cellular toxicity and decreased viability *in vitro*. These results provide clear evidence of toxicity, limiting any potential use of Far-UVC, including 206 nm and 222 nm wavelengths, for pulmonary applications.

## EXPERIMENTAL MODEL AND STUDY PARTICIPANT DETAILS

The studies involving human tissues and informed consent were reviewed and approved by London – Central Research Ethics Committee (REC) (REC No: 16/LO/1883) and by NHS Lothian SAHSC Bioresource (REC No: 20/ES/0061).

## RESOURCE AVAILABILITY

The data presented in this study are available on request from the corresponding author.

## MATERIALS AVAILABILITY

### DATA AND CODE AVAILABILITY

- Microscopy data reported in this paper will be shared by the lead contact upon request.
- Any additional information required to reanalyze the data reported in this paper is available from the lead contact upon request.

## ASSOCIATED CONTENT/SUPPLEMENTARY MATERIAL

### Supplementary figures

**Figure S1.**
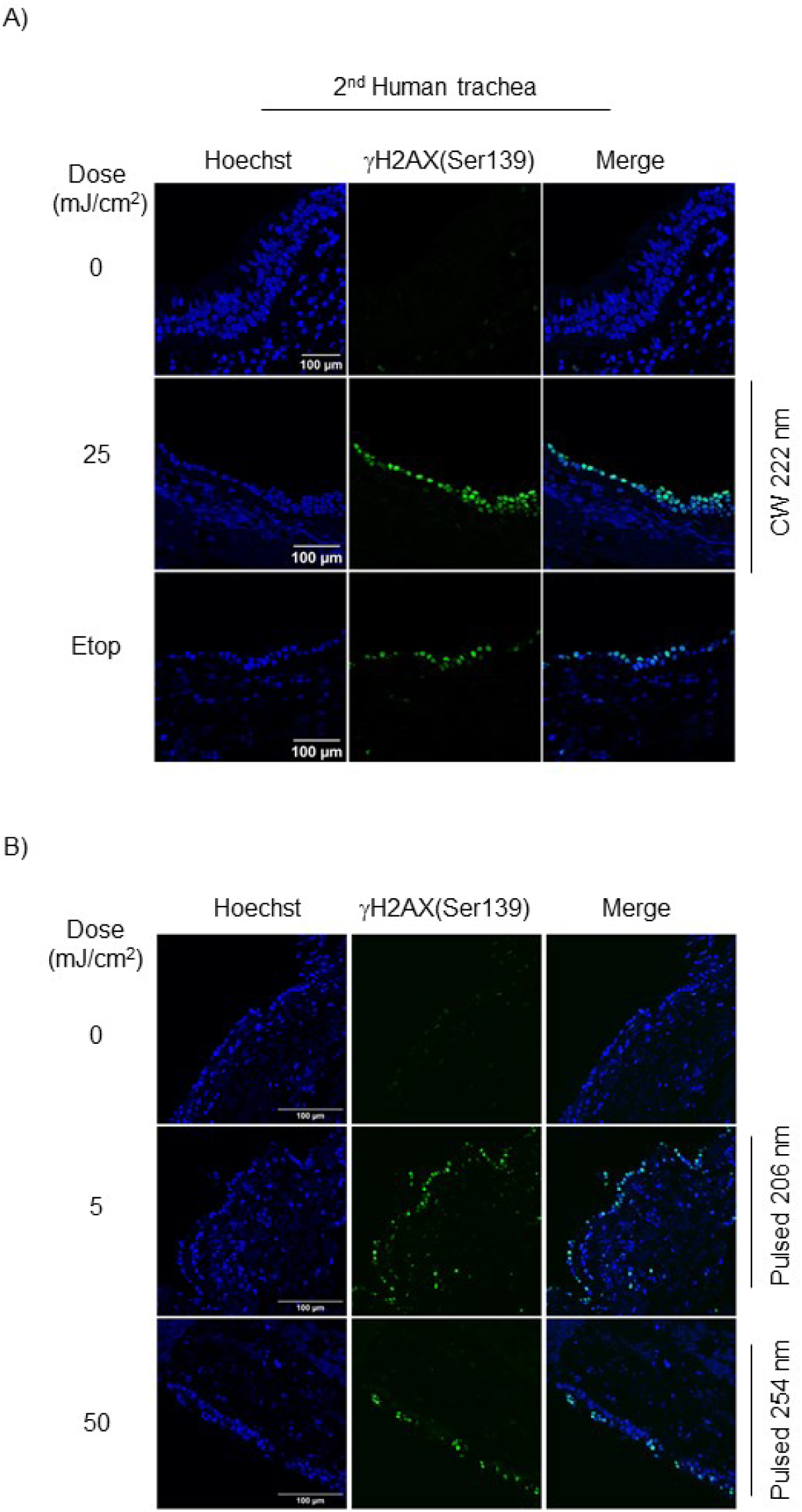
CW 222 nm and pulsed 206 nm irradiation induced the phosphorylation of H2AX(Ser139) in human trachea. (A) Representative immunofluorescence images of γH2AX(Ser139) staining (green) in a second set of human trachea after CW 222 nm irradiation Hoechst nuclei (blue). Scale bar, 100 μm. (B) Representative immunofluorescence images of γH2AX(Ser139) staining (green) after pulsed 206 nm irradiation in human trachea. Hoechst nuclei (blue). Pulsed 254 nm (50 mJ/cm^2^) irradiation was used as a positive control. Scale bar, 100 μm.

**Figure S2.**
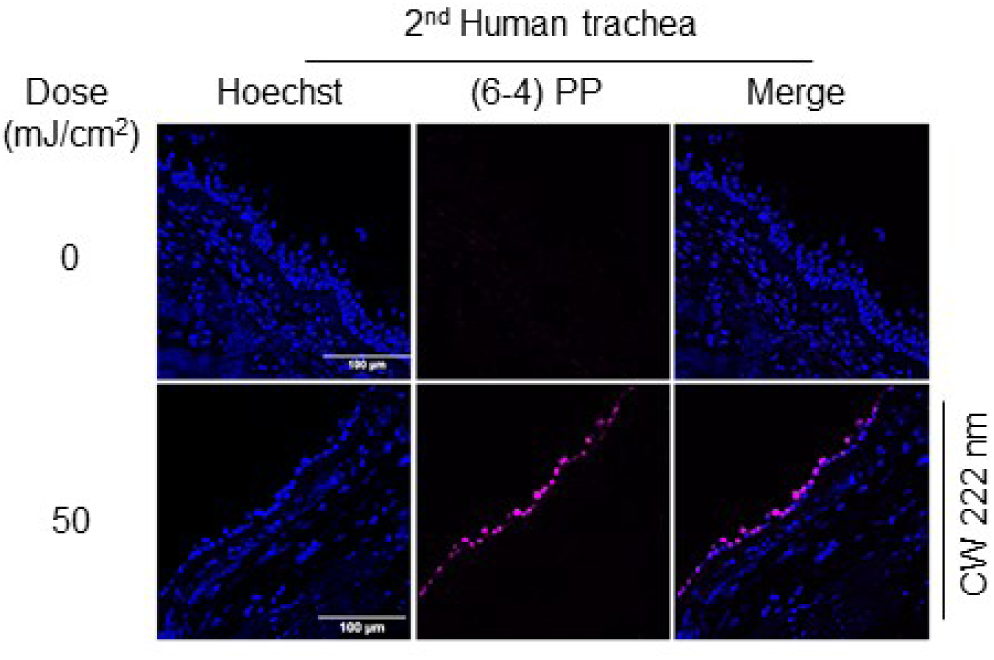
(6-4) PP DNA lesions were detected in CW 222 nm-irradiated human trachea. (6-4) PP staining (magenta) in a second set of human trachea after CW 222 nm irradiation. Hoechst nuclei (blue). Scale bar, 100 μm.

### KEY RESOURCES TABLE

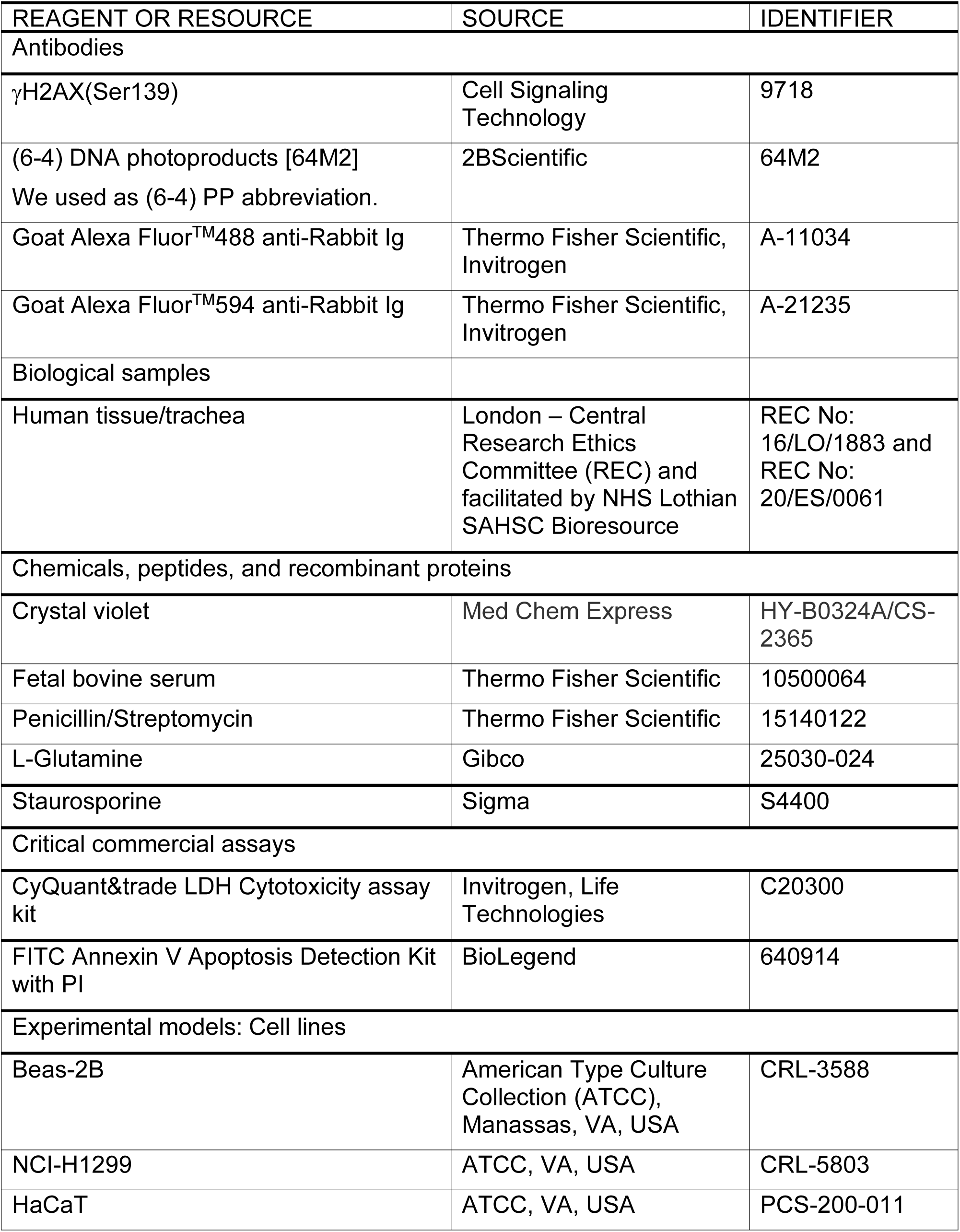
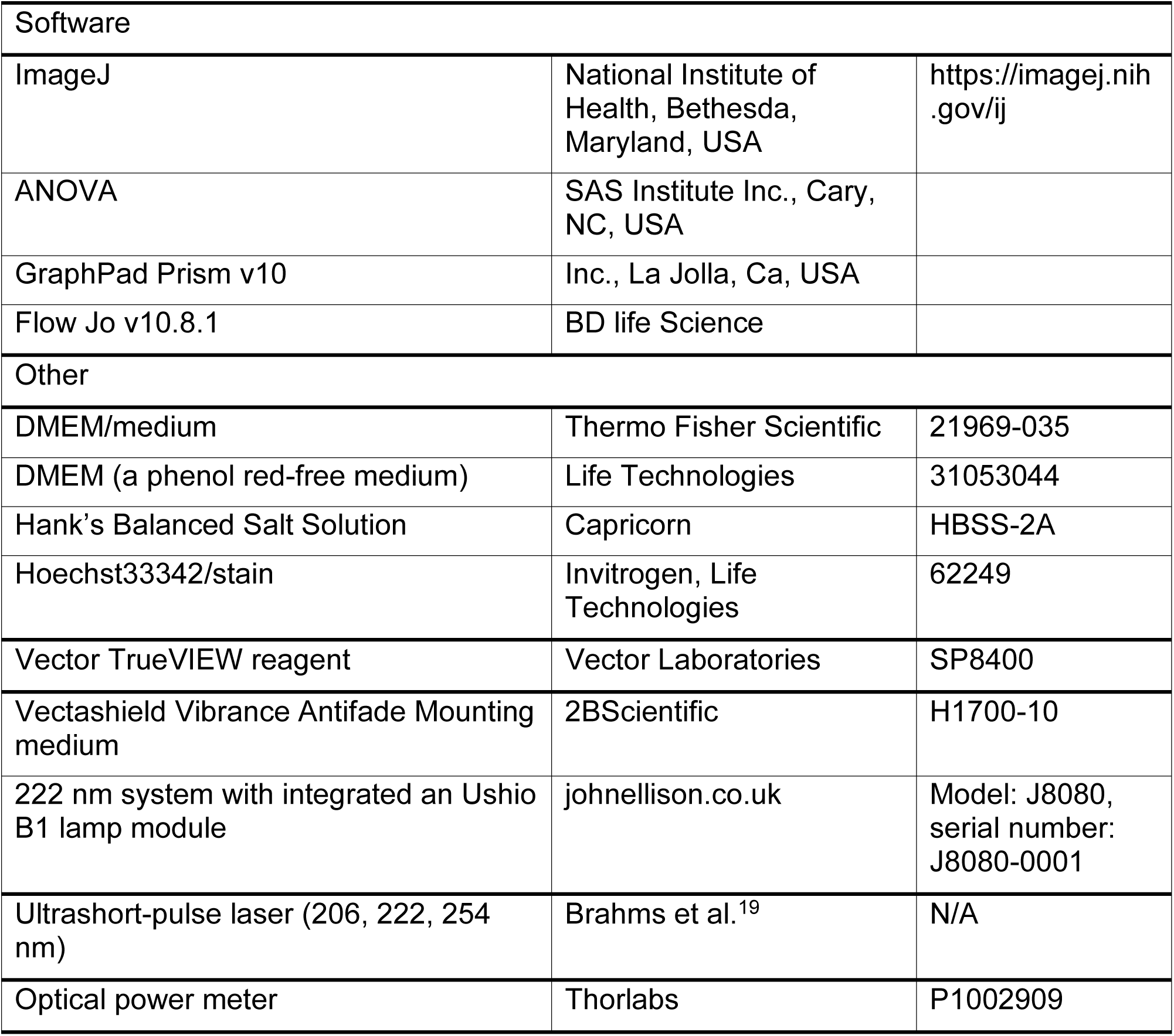

## ACKNOWLEDGMENTS

We thank the QMRI Imaging, IRR SURF Molecular Histology, and IRR Flow Cytometry facilities at the University of Edinburgh for their assistance. Special thanks to Dr Richard O’Connor for help using flow cytometry to study DNA damage response at the beginning of the project. We sincerely thank John Ellison from John Ellison Electronics for kindly providing the 222 nm system integrating an Ushio B1 lamp module. We gratefully acknowledge funding support from the Engineering and Physical Sciences Research Council (EPSRC): EP/T020903/1 and EP/S025987/1, and from the European Research Council under the European Union Horizon 2020 Research and Innovation program: Proof of Concept Grant agreement ULIGHT, No. 899900. C.B. acknowledges support from the Royal Academy of Engineering through Research Fellowship No. RF/202122/21/133. J.C.T. is supported by a Chair in Emerging Technologies from the Royal Academy of Engineering.

## AUTHOR CONTRIBUTIONS

Conceptualization, A.V., J.C.T., R.R.T. and K.D.; methodology, A.V., S.M.P.C.M., C.A.R., M.S., E.W., A.A. and C.B.; experiments, A.V., A.B., J.G., S.M.P.C.M. and C.A.R; writing—review & editing, A.V., N.J., B.M., R.R.T. and K.D.; funding acquisition, A.R.A, L.P, J.C.T, R.R.T and K.D.; resources, S.D., H.X.L., J.G. All co-authors contributed to the preparation of the manuscript, read and approved the submitted version.

## DECLARATION OF INTERESTS

The authors declare no conflict of interest.

## Notes

### Competing Interest Statement

The authors have declared no competing interest.

